# Organ Boundary Circuits Regulate Sox9+ Alveolar Tuft Cells During Post-Pneumonectomy Lung Regeneration

**DOI:** 10.1101/2024.01.07.574469

**Authors:** Tomohiro Obata, Satoshi Mizoguchi, Allison M. Greaney, Taylor Adams, Yifan Yuan, Sophie Edelstein, Katherine L. Leiby, Rachel Rivero, Nuoya Wang, Haram Kim, Junchen Yang, Jonas C. Schupp, David Stitelman, Tomoshi Tsuchiya, Andre Levchenko, Naftali Kaminski, Laura E. Niklason, Micha Sam Brickman Raredon

## Abstract

Tissue homeostasis is controlled by cellular circuits governing cell growth, organization, and differentation. In this study we identify previously undescribed cell-to-cell communication that mediates information flow from mechanosensitive pleural mesothelial cells to alveolar-resident stem-like tuft cells in the lung. We find mesothelial cells to express a combination of mechanotransduction genes and lineage-restricted ligands which makes them uniquely capable of responding to tissue tension and producing paracrine cues acting on parenchymal populations. In parallel, we describe a large population of stem-like alveolar tuft cells that express the endodermal stem cell markers Sox9 and Lgr5 and a receptor profile making them uniquely sensitive to cues produced by pleural Mesothelium. We hypothesized that crosstalk from mesothelial cells to alveolar tuft cells might be central to the regulation of post-penumonectomy lung regeneration. Following pneumonectomy, we find that mesothelial cells display radically altered phenotype and ligand expression, in a pattern that closely tracks with parenchymal epithelial proliferation and alveolar tissue growth. During an initial pro-inflammatory stage of tissue regeneration, Mesothelium promotes epithelial proliferation via WNT ligand secretion, orchestrates an increase in microvascular permeability, and encourages immune extravasation via chemokine secretion. This stage is followed first by a tissue remodeling period, characterized by angiogenesis and BMP pathway sensitization, and then a stable return to homeostasis. Coupled with key changes in parenchymal structure and matrix production, the cumulative effect is a now larger organ including newly-grown, fully-functional tissue parenchyma. This study paints Mesothelial cells as a key orchestrating cell type that defines the boundary of the lung and exerts critical influence over the tissue-level signaling state regulating resident stem cell populations. The cellular circuits unearthed here suggest that human lung regeneration might be inducible through well-engineered approaches targeting the induction of tissue regeneration and safe return to homeostasis.

## Introduction

Pulmonary regenerative medicine requires an understanding of tissue-level control mechanisms that govern lung homeostasis. Single-cell atlases have been used extensively to study pulmonary populations, but comparatively little study has been done of the extracellular circuits controlling dynamic tissue remodeling. In this study, we identify previously undescribed cellular circuits that control post-pneu-monectomy lung regeneration. Our results demonstrate that pleural mesothelial cells coordinate a global, tissue-level response to increased tissue tension by modulating their expression of morphogenic ligands sensed by parenchymal populations and, in particular, Sox9+/Lgr5+ alveolar tuft cells.

Post-pneumonectomy lung regeneration is a dynamic process in which lung tissue grows new airways, vasculature, and gas-exchanging alveoli following surgical lobectomy (*1-5*). This process is known to be dependent on mechanical tissue tension in the residual lung tissue, and is thought to be driven by a combination of YAP/TAZ-mediated HIPPO-family cell-signaling cascades in combination with immune cell recruitment into the tissue (*3, 6, 7*). However, the exact cellular sensors and effectors involed in this process are currently undefined. Post-pneumonectomy lung growth is a unique and powerful context in which to study tissue regeneration dynamics, because cellular turnover increases in the absence of direct tissue injury or infection (*8*). In rats, residual lung tissue displays altered histology by day 3 post-lobectomy, expands rapidly around day 7, and returns to homeostasis during days 14-21 (*9, 10*). There is not scientific consensus on how well this rodent process mirrors human *in vivo* biology, largely due to the difficulty of obtaining reliable data which would capture this process in human beings. However, it is clear that rats have a remarkable capacity for healthy, well-regulated alveolar regeneration which could inform the development of regenerative therapeutics.

It has been known for almost 70 years that the lungs of rats contain abundant chemosensitive alveolar tuft cells that are comparatively rare in the lungs of both mice and humans (*11-13*). Alveolar tuft cells have been given many names since their first discovery in 1956, including brush cells, caveolated cells, multivesicular cells, fibrillovesicular cells (*12, 14*), solitary chemosensory cells (*15, 16*), and even alveolar type III pneumocytes (*14, 17-19*) due to their direct contact with alveolar type I (ATI) and alveolar type II (ATII) cells in the distal alveoli. Their biological function in the alveoli has been discussed with great interest over the decades but is still currently unknown (*12, 14, 20, 21*). Tuft cells, as a general cell type, are found throughout the epithelium of many tissues, including the respiratory tract, liver, pancreas, thymus, and gastrointestinal tract. They are marked, across tissue, by expression of the taste-receptor gene *Trpm5*, the transcription factor *Pou2f3*, and the cell-plasticity-associated transcript *Dclk1*. In the lung, tuft cells are thought to play a sentinel-like chemosensory role for inhaled pollutants (*14, 19, 21-27*). In the trachea and bronchi, tuft cells communicate with afferent nerve fibers to mediate central physiologic responses such as coughing (*16, 22, 28-30*). In the gut, tuft cells influence mucosal and innate immunity (*22, 28, 31*). In the pulmonary alveoli, however, the function of tuft cells is not well defined.

We show in this manuscript that alveolar tuft cells in the rat strongly express the canonical stem-cell markers *Sox9* and *Lgr5*, during both tissue homeostasis and during post-pneumonectomy lung regeneration. *Sox9* is a master regulator of cellular plasticity across many endodermal and neural-crest-derived tissues during both development and adult homeostasis. *Sox9* marks multipotent populations in the lung, gut, liver, pancreas, prostate, kidney, thymus, testes, cartilage, and retina (*32, 33*), and its prominence in the alveolar tuft cell transcriptome strongly suggests that these cells may fulfill a progenitor function. *Lgr5* is a canonical stem-cell-surface receptor which transduces paracrine WNT cues. In the gut, joint *Sox9* and *Lgr5* expression marks a distinct population of reserve stems cells which proliferate only in response to sever tissue injury or when high epithelial turnover is required (*34-36*). The display of this specific regenerative signature in alveolar tuft cells suggested to us the likely presence of extracellular circuits either constraining or controlling alveolar tuft cell progenitor behavior. We therefore created a dynamic atlas of extracellular signaling during post-pneumonectomy lung regeneration, and we show in this manuscript that alveolar tuft cells likely respond to morphogenic cues originating from the pleural mesothelium.

## Results

### Alveolar Tuft Cells Prominently Express Sox9 and Lgr5

We created a homeostatic adult rat lung atlas by integrating scRNAseq data from 8 male and 6 female rat tissues dissociated via intravascular enzyme perfusion and low-shear suspension (Fig 1A, see Methods). The resulting dataset contained 20,225 immune cells (49.2%) 11,699 epithelial cells (28.4%), 6,365 endothelial cells (15.5%), and 2,854 mesenchymal cells (6.9%) (Fig 1B). These four cellular classes were clustered into 34 distinct cell types for downstream analysis which can be seen in Fig 1C-F. Within epithelium, we identified ciliated cells, secretory cells, ATI cells, ATII cells, ATII-ATI transitional cells, bronchio-alveolar stem cells (BASCs) and a prominent population of tuft cells (Fig 1C). Tuft cells were the third most common epithelial cell type captured, comprising 8.8% of all captured epithelium and 2.5% of the total tissue, after ATII cells (38.8%; 11.0%) and ATI cells (37.0%; 10.5%). BASCs were the least common epithelial cell type, at 1.7% of epithelium and 0.48% of the whole tissue. In alignment with previous reports from mice and humans, we identified five major endothelial subpopulations, annotated as general microvascular endothelial cells (gCaps), aerocytes (aCaps), and venous, arterial, and lymphatic endothelial cells (Fig 1D). Mesenchymal cells were divided into seven distinct populations, annotated as *Col13a1*+ fibroblasts, *Col14a1*+ fibroblasts, *Lgr5*+ fibroblasts, mesothelium, pericytes, smooth muscle cells, and myofibroblasts (Fig 1E). The immune cells were clustered into 18 populations, annotated as B cells, basophils, cycling immune, dendritic cells, eosinophils, innate lymphoid cells, alveolar macrophages, proliferating alveolar macrophages, interstitial macrophages, mast cells, activated monocytes, classical monocytes, nonclassical monocytes, neutrophils, natural killer cells, plasma cells, T cells, and killer T cells (Fig 1F).

**Figure 1:**
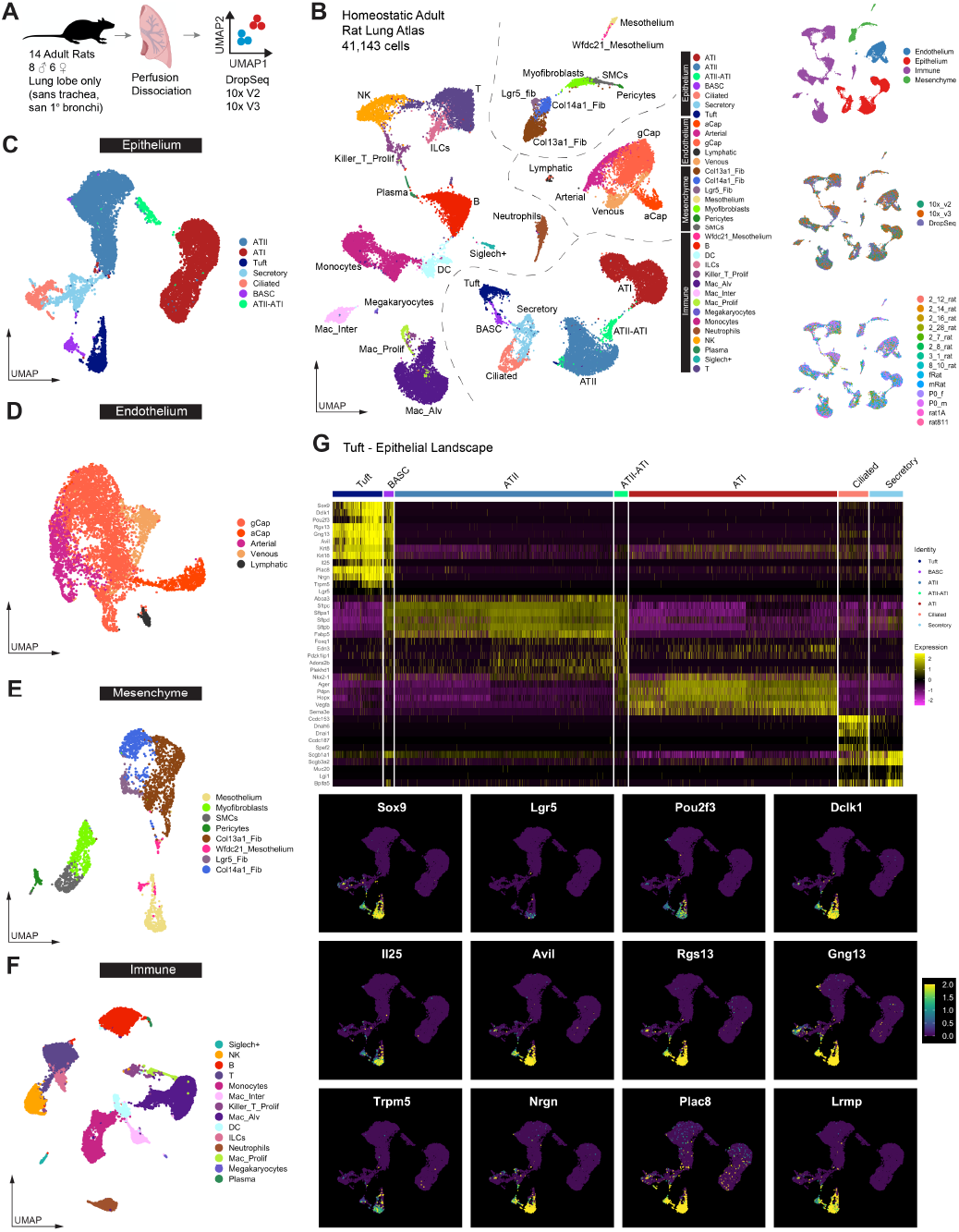
Homeostatic adult rat lungs contain abundant tuft cells expressing Sox9 and Lgr5. **(A)** Schematic of single-cell data acquisition. **(B)** Homeostatic rat lung atlas containing 41,143 cells grouped into four cell classes and 34 cell types. **(C)** Epithelial cells include ATII, ATI, Tuft, Secretory, Ciliated, ATII-ATI Transitional, and BASCs. **(D)** Endo-thelial cells include gCaps, aCaps, Arterial, Venous, and Lymphatic. **(E)** Mesenchymal cells include Mesothelium, Myofibroblasts, Smooth Muscle Cells, Pericytes, Col13+, Col14+, and Lgr5+ Fibroblasts, and a small population of Mesothelial cells positive for *Wfdc21*. **(F)** Immune cells include Alveolar and Interstitial Macrophages, B cells, T cells, Monocytes, Natural Killer cells, Plasma cells, Megakaryocytes, Neutrophils, Dendritic cells, Killer-T cells, and Innate Lymphoid cells. **(G)** Heatmap showing selected markers defining the epithelial cell landscape. Tuft cells prominently express *Sox9* and *Lgr5*, along with canonical tuft markers including *Pou2f3, Dckl1, Avil, Trpm5, Il25, Rgs13* and *Ggn13*. BASCs express lower values of all tuft cell genes, but in combination with distinct and unique co-expression of ATII markers, including *Sftpa, Sftpb, Sftpc, Sftpd* and *Abca3*, as well as club cell markers including *Scgb1a1* and *Scgb3a2*.

Investigation of epithelial phenotype showed that Tuft cells and BASCs share transcriptomic character (Fig 1G). Both populations uniquely express stem-like genes *Dclk1, Sox9* and *Lgr5*, with tuft cells expressing these genes at distinctly higher levels. BASCs were marked by additional expression of club-cell marker *Scgb1a1* and ATII markers *Abca3* and *Sftpc*, which were not expressed by tuft cells. Tuft cells were marked by additional expression of taste-receptor *Trpm5* and transcription factor *Pou2f3*. Tuft cells can be clearly identified from other cells in the lung as being *Dclk1*-positive/*Scgb1a1-*negative or *Sox9*-positive/*Abca3*-negative. These signatures were used to identify Tuft cells in subsequent analyses.

Immunofluorescent staining showed BASCs to reside exclusively in the bronchioalveolar junctions (Fig 2A-B). In contrast, tuft cells were common throughout the alveoli (Fig 2C-F). Alveolar tuft cells prominently expressed *Sox9* and were clearly distinguishable from both ATI (Fig 2F) and ATII cells (Fig 2G-I). We found tuft cell markers and ATII markers overlapped very rarely in the alveolus (Fig G-I). Together with the scRNAseq data presented in Figure 1, these findings confirmed alveolar tuft cells to be a distinct and common resident epithelial cell type in the rat alveolus. Their unique expression of known stem cell markers led us to hypothesize that alveolar Tuft cells may serve an important regenerative role unfulfilled by other alveolar-resident epithelial cell types.

**Figure 2:**
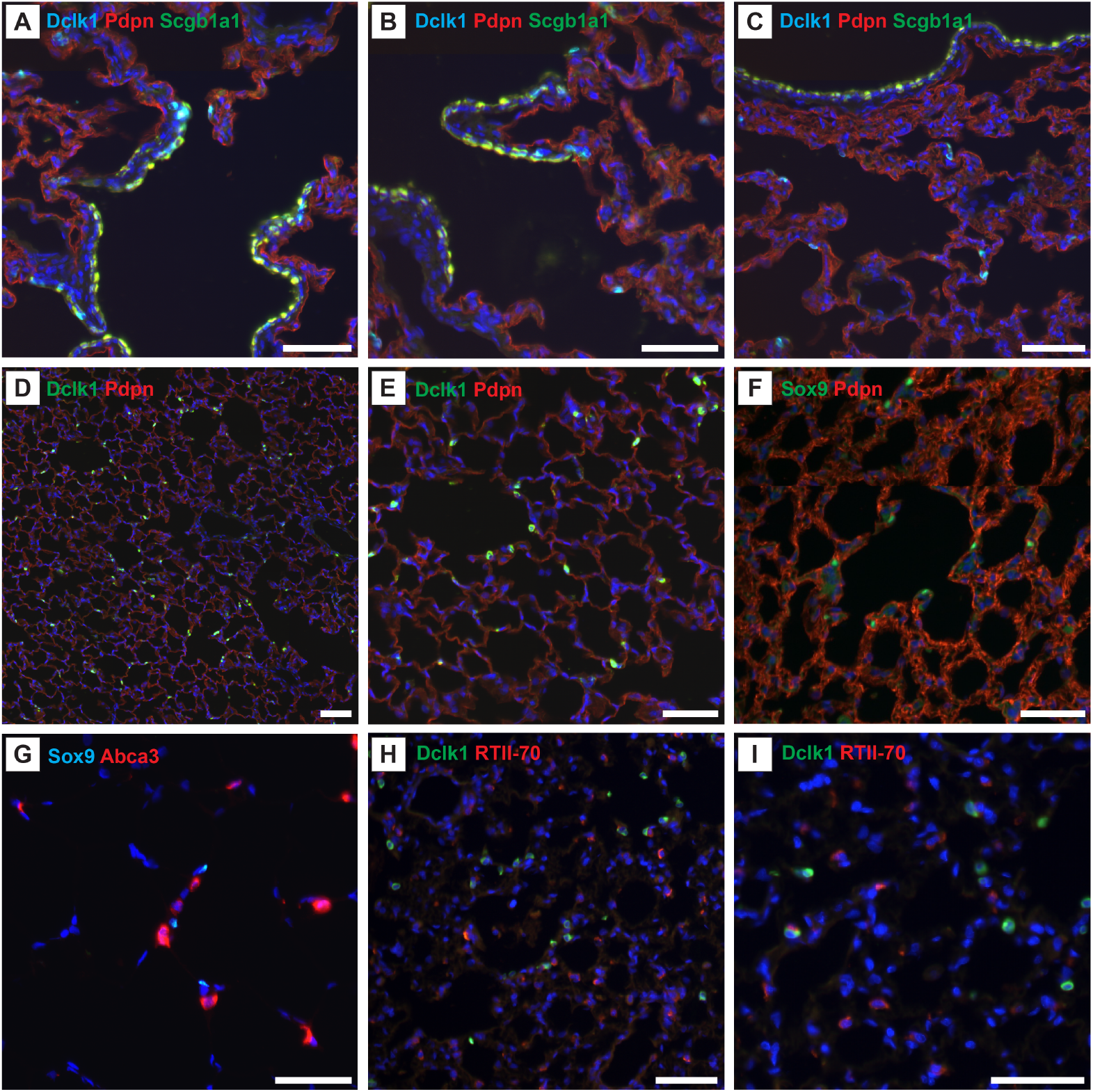
*Sox9*+ Tuft Cells are present throughout the alveoli of adult rats. **(A-C)** Immunostaining of the bronchioalveolar junction and airway-adjacent alveolar tissue. The bronchioalveolar junction consistently displays *Dclk1*+/*Scgb1a1*+ BASCs, while more distal alveolar regions contain abundant *Dclk1*+/*Scgb1a1*-tuft cells. *Pdpn* shows ATI cell localization. **(D-E)** *Dclk1*+ tuft cells are extremely common in the rat alveolus and almost invariably contact ATI cells. **(F)** Alveolar tuft cells uniquely express the transcription factor *Sox9*. **(G-I)** *Sox9*+/*Dclk1*+ alveolar tuft cells are distinct from ATII cells. Scale bars are 50μm.

### Alveolar Tuft Cells are Specifically Tuned to Extracellular WNT Morphogens Expressed by Pleural Mesothelial Cells

We next analyzed patterns of ligand-receptor connectivity in the homeostatic lung, using the software package NICHES (*37*). NICHES operates by randomly sampling 1:1 pairings of cells within a dataset (Fig 3A) to create a comprehensive, quantitative atlas of cell-to-cell connectivity within a tissue. The resulting data can be used in two powerful and distinct ways: first, to create a single-cell-barcoded proxy of cellular microenvironment (Fig 3B-C) and second, to embed, cluster, and directly analyze groupings of individual cell-to-cell signaling edges present within tissues (Fig D-F). Our analysis yielded two main conclusions important for this study.

**Figure 3:**
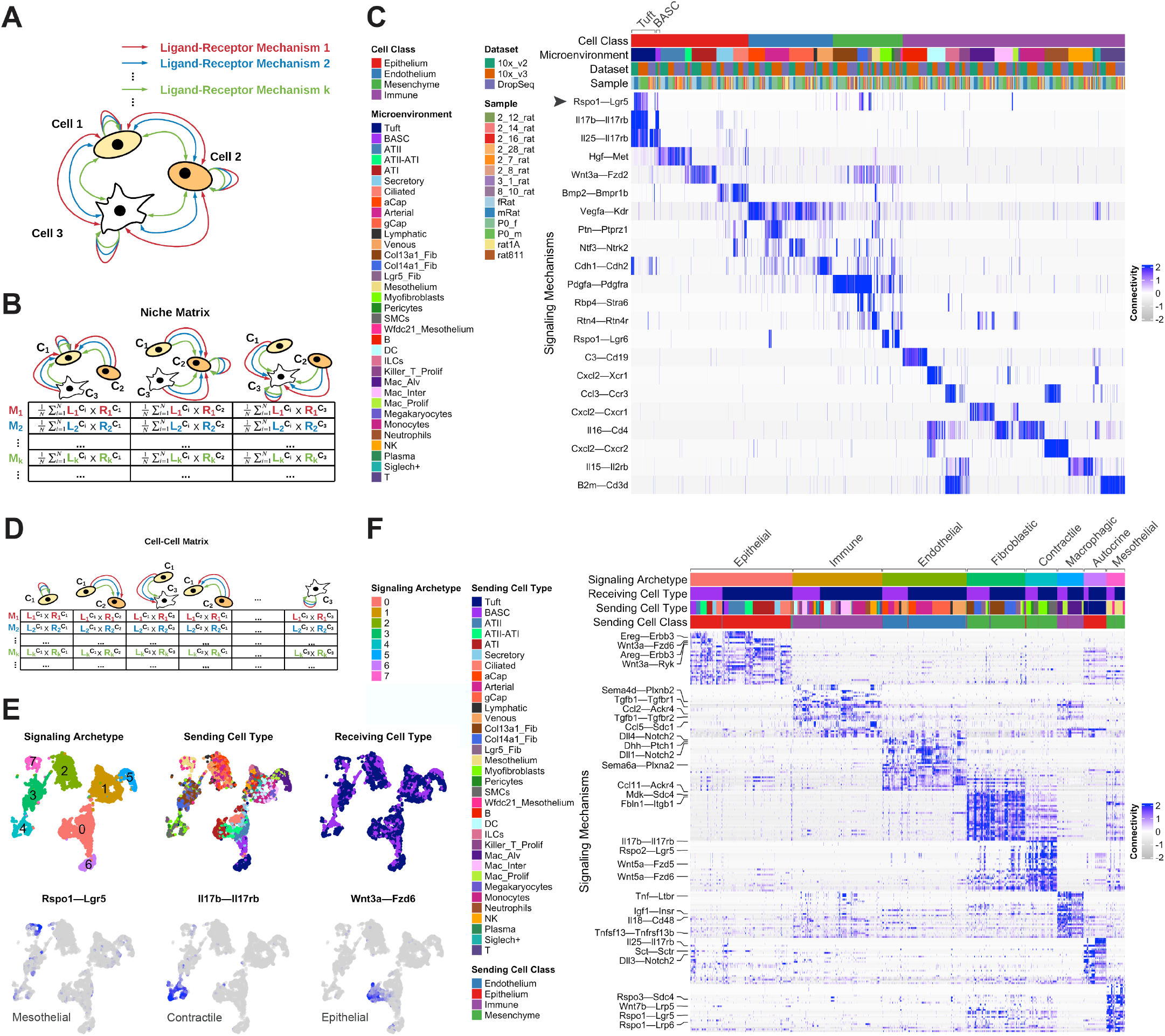
Alveolar tuft cells and BASCs are uniquely tuned to extracellular WNT-morphogen signaling from mechanosensitive cells, including mesothelial cells. **(A)** Schema of cells interacting via multiple ligand-receptor mechanisms. **(B)** Schematic of System-To-Cell NICHES data, which can be analyzed to reveal signaling mechanisms specific to particular cellular niches within a tissue. **(C)** Heatmap of signaling connectivity within homeostatic lung tissue. Note the specificity of *Rspo1*—*Lgr5*, marked with a black arrow, as well as *Il17b*—*Il17rb* and *Il25*—*Il17rb* to the tuft cell and BASC microenvironments. Note also that ATII cells are uniquely receptive to signaling via *Hgf*—*Met*, ATI cells to signaling via *Wnt3a*—*Fzd2*, and proximal epithelium to signaling via *Bmp2*-*Bmpr1b*. **(D)** Schematic of Cell-To-Cell NICHES data. **(E)** Louvain clustering of cell-to-cell signaling edges landing on Tuft cells and BASCs within the lung. We identified eight distinct archetypes (top-left). Metadata showing sending and receiving cell type for each edge is included (top-middle and top-eight), as well as feature-plots of selected connectivity mechanisms of interest. Labeling of each signaling archetype based on sending cell category revealed distinct networks acting on tuft cells associated with epithelial, immune, endothelial, fibroblastic, contractile, macrophagic, autocrine, and mesothelial cell sources. **(F)** Heatmap showing marker mechanisms enriched within each signaling archetype. The mesothelial-tuft relationship has unique character, distinct from all other relationships within the lung, and is marked prominently by stem-modulating WNT-morphogens including *Rspo1—Lgr5, Wnt7b—Lrp5*, and *Rspo3—Sdc4*. Note also the presence of *Wnt3a—Fzd6* connectivity from ATI cells and *Wnt5a—Fzd5/6* and *Il17b—Il17rb* connectivity from contractile mesenchymal cells.

First, we found that alveolar tuft cells and BASCs were uniquely tuned to major WNT-family signaling ligands produced by other populations in the lung. Key mechanisms of biological interest included *Rspo1, Il17*, and *Il25* (Fig 3C). Our data showed that signals sent via these mechanisms would be most likely, by far, to be sensed by alveolar tuft cells rather than other parenchymal cells.

Second, we found that mesothelial-epithelial edges cluster separately from others within the Tuft/BASC niche (Fig 3E, cluster 7 and Fig 3F, ‘Mesothelial’). This cluster is marked distinctly by connectivity via *Rspo1—Lgr5/Lrp6, Wnt7b— Lrp5*, and *Rspo3—Sdc4*, all of which are morphogenic mechanisms known to regulate stem cell growth and spatial patterning. The unique mesothelial-tuft connectivity by these specific mechanisms led us to hypothesize that maybe we were observing evidence of a sensor-effector loop regulating organ size. Because mesothelial cells exist solely on the outside surface of the lung, they are in a unique position to monitor tissue strain and to coordinate a parenchymal response. We decided to test this hypothesis by closely studying the role of both alveolar tuft cells and pleural mesothelial cells in post-pneumonectomy lung regeneration.

### Alveolar Tuft Cells Proliferate and Differentiate following Pneumonectomy

We first optimized a survival-surgery protocol to sterilly remove the left lung from adult rats (Fig 4A). This allowed us to carefully observe the response of the remaining tissue to this surgery using time-course whole-slide histology. During the experimental period, the remaining tissue roughly doubled in size (Fig 4B), with tissue regeneration progressing ‘from the outside in’ within affected lobes (Fig 4B-C, data from accessory lobe shown). The first observe histologic change was a thickening and darkening of mesothelial layer, which immediately preceded immune cell recruitment and local parenchymal remodeling. The resolution of this process also follows an outside-in pattern, with the subpleural regions returning to alveolar homeostasis in advance of central regions farther from the mesothelial layer (see Fig 4B, right-most panel).

**Figure 4:**
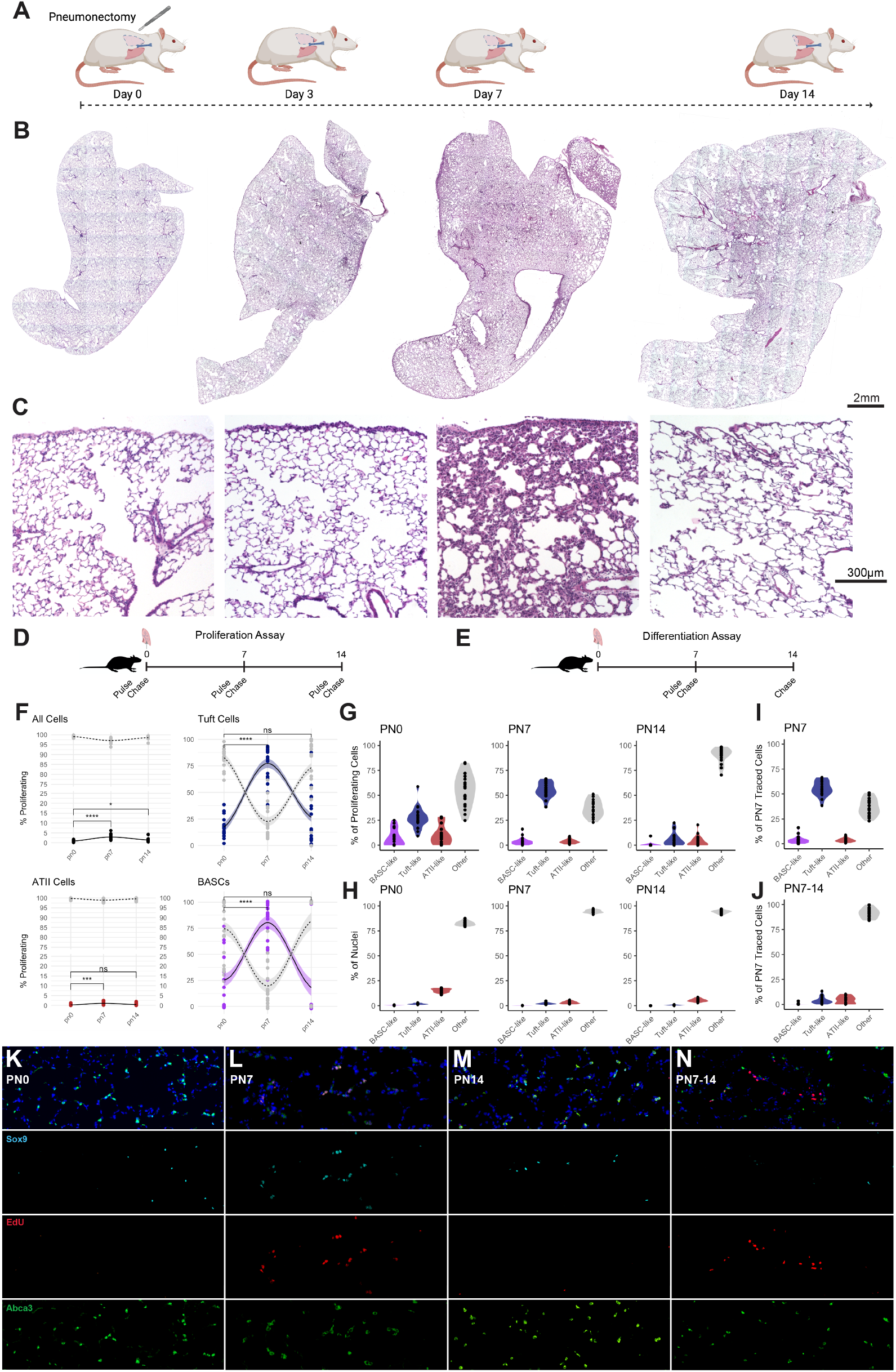
Post-pneumonectomy lung growth triggers proliferation and differentiation of Sox9+ alveolar Tuft Cells. **(A)** Schematic of surgical approach. **(B)** Whole-slide scale-matched histology of the remaining lung tissue at post-surgical day 0, 3, 7, and 14. Note the ‘outside-in’ pattern of regenerative inflammation. **(C)** Scale-matched histology of the subpleural region of the lung over the same time-course. Note the thickening of the pleural border at day 3, widespread alveologenesis at day 7, and return to homeostasis by day 14. **(D)** Schematic of experiment to assess cellular proliferation. **(E)** Schematic of experiment to assess cellular differentiation. **(F)** Fraction of all cells, tuft cells, ATII cell, and BASCs proliferating at selected timepoints. **(G)** Cell type contribution to proliferating cell pool, expressed as fraction of total proliferating cells. **(H)** Cell type contribution to total cell pool, expressed as fraction of nuclei. **(I)** Composition of proliferating cell pool at day 7 post-pneumonectomy. **(J)** Composition of the same traced pool at day 14. **(K)** Histology of tissue pulsed at day 0, **(L)** day 7, **(M)** and day 14. **(N)** Histology of tissue pulsed at day 7 and harvested at day 14. * p < 0.05, ** p < 0.01, *** p < 0.001, **** p < 0.0001

We then inquired which epithelial cell types in the alveolus displayed the greatest proliferation following pneumonectomy. To test this, we performed *in vivo* pulse-chase labeling experiments with 5-ethynyl-2’-deoxyuridine (EdU) during homeostasis and at post-surgical days 3, 7, and 14 (Fig 4D,E). EdU is incorporated exclusively into the DNA of proliferating cells and has a short half-life allowing precise pulse-chase experiments (*38*). These experiments found that during homeostasis only ∼10% of tuft cells and ∼25% of BASCs are actively proliferating (Fig 4F). However, following pneumonectomy, upwards of 75% of both the tuft cell and BASC population became highly proliferative and displayed extensive EdU uptake (Fig 4F).

These same data were then processed to determine which cells within the entire tissue, epithelial or not, proliferated over time (Fig 4G). During homeostasis, tuft cells represent nearly 25% of all proliferating cells in the rat lung (Fig 4G, left), even though they represent only a small fraction of the total nuclei surveyed (Fig 4H, left). By day 7 following pneumonectomy, over 50% of the total proliferating cells in the tissue are tuft cells (Fig 4G, middle). Interestingly, however, this change is associated with minimal shift in the ratio of different cell types to one another (Fig 4H, middle). By day 14 post-pneumonectomy, ∼90% of proliferating cells are non-epithelial (Fig 4G, right), even though the tuft-population proliferation rate did not drop below normal homeostatic levels (Fig 4F, top-right). Again, we observed minimal change in cell type ratios associated with altered cellular proliferation. This observation, namely, that altered cellular proliferation was not associated with clear changes in relative population ratios, strongly suggested to us the presence of robust systems-level regulatory mechanisms controlling cell type ratios. The existence of cellular circuits controlling this behavior would be in close alignment with seminal work showing the importance of cell-cell signaling in the regulation of population dynamics during tissue change (*39*).

Finally we asked if we could observe evidence to suggest tuft-cell differentiation into other cell types as the tissue returns to homeostasis. Affirmative data would reinforce our developing hypothesis that alveolar tuft cells were acting as an alveolar progenitor during post-pneumonectomy lung regeneration. To query this, we performed a longitudinal pulse-chase study (Fig 4D), comparing the character of proliferating cells at day 7 to the character of this identical labeled population one week later at day 14. At day 7, 50% of all proliferating cells were tuft cells (Fig 4I). However, by day 14, less than 10% of this same traced population still displayed tuft-cell character (Fig 4J). Instead, we observed population-scale adoption of a cellular state displaying neither Tuft, BASC, or ATII markers (Fig 4J). This suggested to us that either a substanital fraction of the tuft cells proliferating at day 7 were differentating into ATI cells or, possibly, were undergoing unobserved epithelial-to-mesenchymal transition, which would look similar in these data.

### Adult Dclk1+ Cells Rapidly Produce Sox9+ Organoids

To more deeply probe tuft cell multipotency in a controlled *in vitro* model, we developed an organoid protocol to grow epithelial cells isolated from the distal adult lung using a magnetic bead pull-down based on protein expression of *Dclk1* (see Methods and Figure S1). This surface marker (*40*) was identified from our single-cell data as highly specific to tuft cells and BASCs jointly. The resulting single-cell suspension was 78.75% ± 1.98% Sox9+ (n=45), 31.33% ± 3.41% BASCs (Sox9+/Scgb1a1+) and 68.67% ± 3.41% Tuft Cells (Sox9+/Scgb1a1-). This distribution aligned closely with our *in situ* homeostatic single-cell data, demonstrating the desired unbiased capture of both *Dclk1*-positive epithelial populations and minimal non-epithelial contaminants.

The isolated cells expanded rapidly in matrigel into three-dimensional spherical organoids (Fig 5A-B). We adapted a protocol that was first pioneered to grow *Sox9*-positive progenitors from embryonic mouse lungs, which uses a combination of WNT-agonism, ROCK-inhibition, TGFB-inhibition, p38 MAPK-inhibition, and FGF- and EGF-agonism (*41*). We found that organoid growth was improved by substituting rat, rather than murine, growth factors (data not shown), and all further experiments were performed with this rat-adapted protocol (see Methods). After 7 days of growth, these organoids displayed distinctly BASC-like character, co-expressing Sox9, CCSP, Abca3, lowly expressing proSPC and RTII-70, and notably losing all observable tuft cell character including expression of Dclk1 (Fig 5C-K). Additionally, these organoids proved to be negative for key non-BASC markers, including Pdpn (ATI cells), Krt5 (basal cells), Mucin-5AC (secretory cells) and alpha-Tubulin (ciliated cells). They were highly proliferative and could be passaged repeatedly without observed change in character for many months.

**Figure 5:**
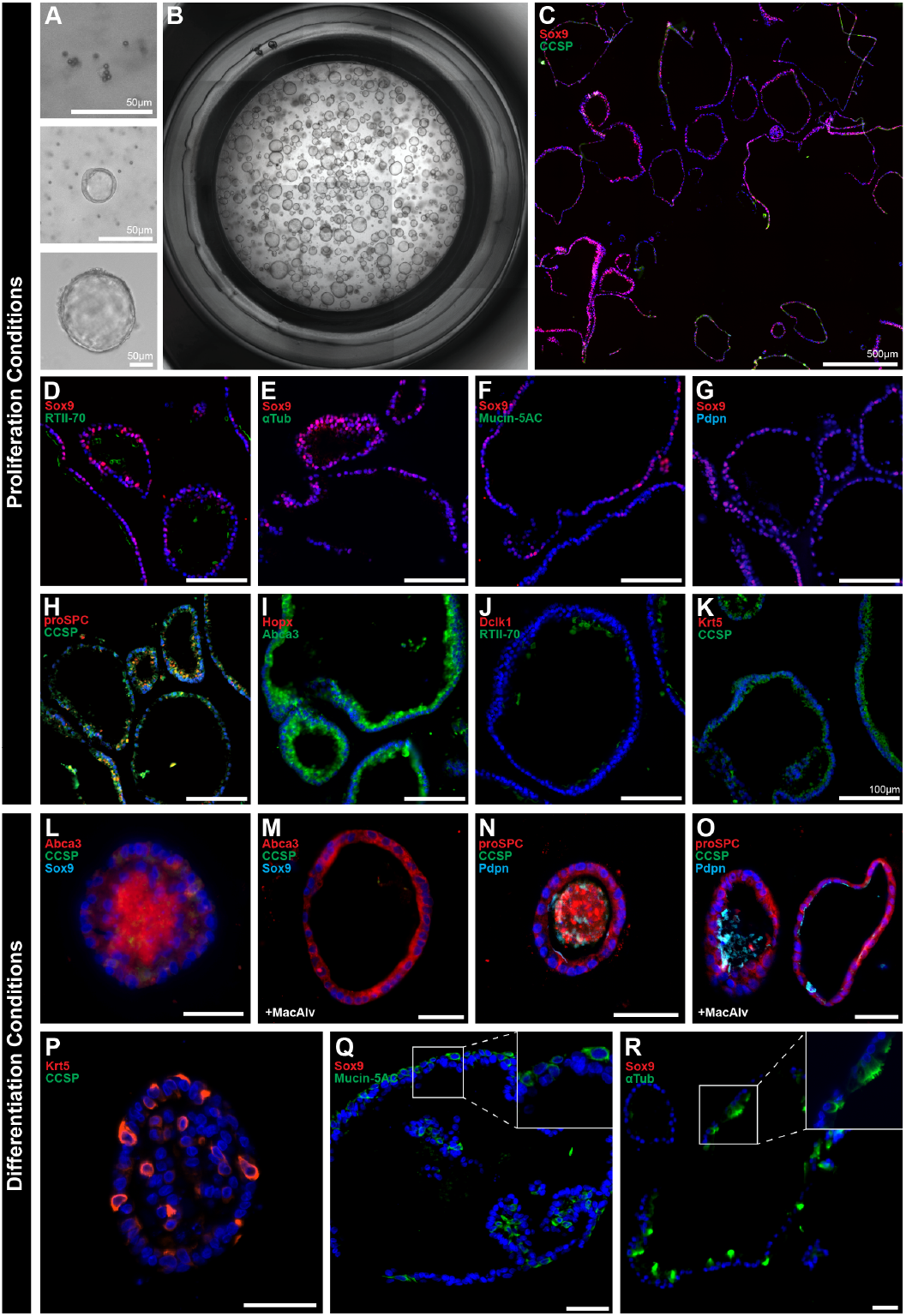
Dclk1+ epithelial cells yield highly proliferative Sox9+ or-ganoids that stochastically differentiate into both proximal and distal lineages. **(A-B)** Light microscopy of organoid growth over time. In proliferation media, Dclk1+ cells grew rapidly in 3D culture and developed into homogenous hollow spherical organoids. **(C)** Immunostaining of day 7 organoids for Sox9 and Scgb1a1 (CCSP). Note near-universal organoid positivity for Sox9. **(D-K)** Immunostaining for selected markers. We found many organoids to display a BASC-like phenotype, co-expressing Sox9 concurrently with RTII-70, Scgb1a1, Sftpc, and Abca3, while not expressing terminal differentiation markers such as acetylated tubulin, Mucin5AC, Pdpn, Hopx, Dclk1, or Krt5. **(L-R)** Immunohistochemistry of organoids grown in differentiation conditions, stained for selected lineage markers. When transitioned to differentiation media, the organoids reduced their expression of Sox9 and Scgb1a1 and began stochastic differentiation toward diverse distal-like **(L-O)** and proximal-like **(P-R)** cell states. The addition of alveolar macrophages (see **M** and **O**) reduced the presence of intra-organoid debris and promoted expression of the ATII marker protein Abca3 and the ATI marker protein Pdpn.

### Sox9+ Organoids Display Stochastic, Milieu-Dependent Differentiation into both Proximal and Distal Pulmonary Lineages

We then tested the ability of the cells within these organoids to differentiate. After 7 days of organoid expansion, we removed all of the above small molecules and growth factors from the media except except EGF, modifying a protocol from (*42*). The organoids changed substantially following this transition, becoming heterogeneous in morphology within days and displaying stochastic, organoid-specific expression of both proximal and distal lung lineages after two weeks of culture (Fig 5L-R). Organoids could be found on immunostaining displaying marker profiles of all known rat lung lineages, including ATI cells (Pdpn+), ATII cells (Abca3+/Sox9-), Club cells (Scgb1a1+/Sox9-), Ciliated cells (aTub+), goblet cells (Muc5ac+) and basal cells (Krt5+). The addition of alveolar macrophages increased clearance of intraluminal debris, supported stem-cell character and organoid growth and promoted distal differentiation (Fig 5L-O). Notably, very few organoids within differentiation conditions retained BASC-like character (Sox9+/Scgb1a1+/Abca3+) without the addition of macrophages. We hypothesized from this observation that extracellular WNT-family signaling cues were likely enforcing the stem-like character of Sox9+ alveolar tuft cells *in vivo*, and that this signaling family in particular might hold clues to the regulation of tuft cell stemness *in vivo*. We decided to test this hypothesis directly by comprehensively mapping the changes in cell-signaling architecture occuring during post-pnuemonectomy lung growth and testing which pathways were most intensely perturbed.

### Post-Pneumonectomy Lung Growth Reveals Regulatory Circuits Influencing Progenitor Function of Alveolar Tuft Cells

We generated single-cell data from the accessory lobe of both male and female rats in parallel from post-surgical Day 0, 3, 7, and 14 (see Methods). These data were sequenced at a common read depth, cleaned, annotated, tagged with appropriate metadata, and assessed without integration. An overview of the resulting cell atlas can be seen in Fig 6A-B.

**Figure 6:**
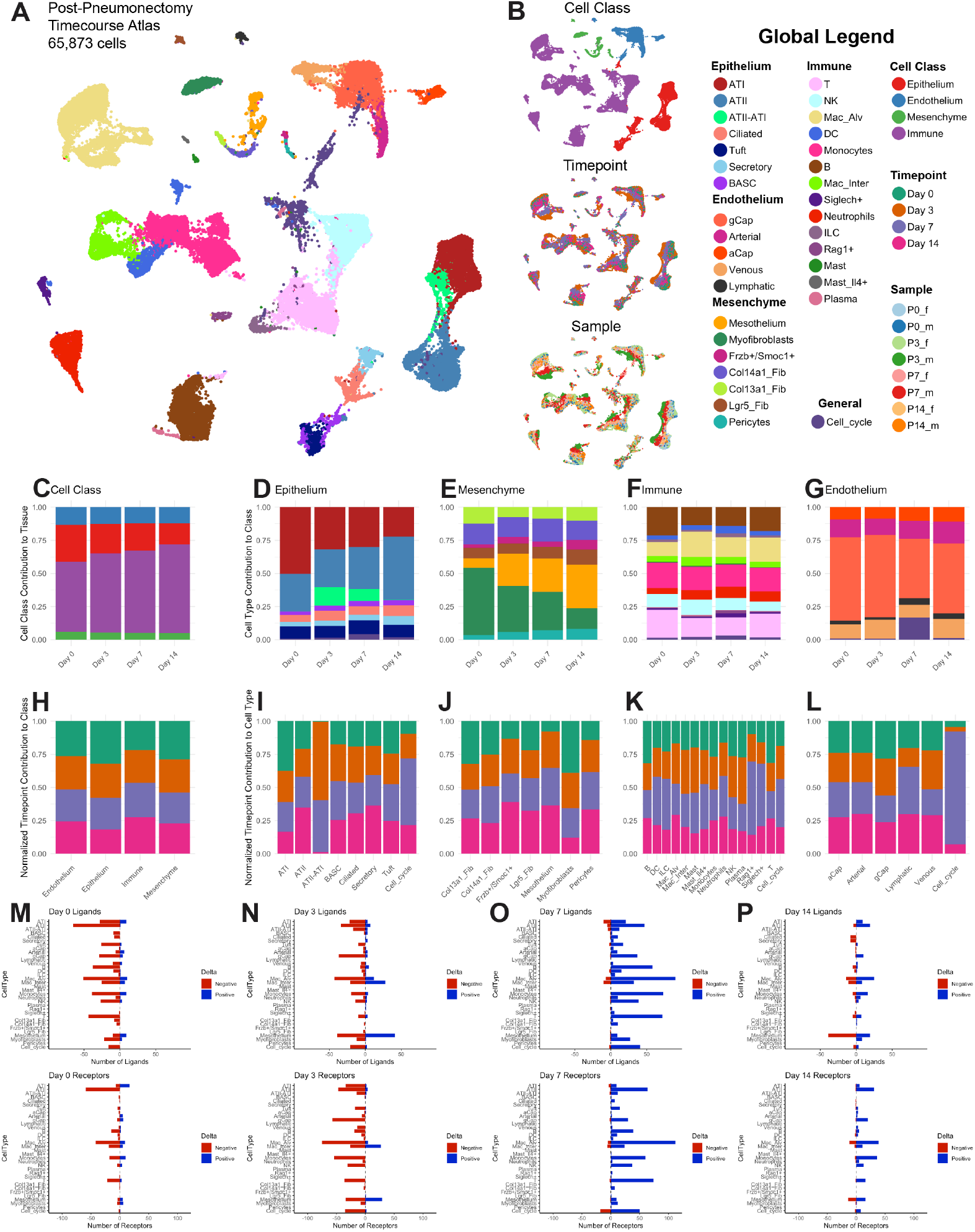
Increased epithelial flux is associated with altered ligand-receptor connectivity. **(A)** Post-pneumonectomy time-course atlas, embedded without data integration. **(B)** Class, timepoint, and sample metadata distrobutions. **(C)** Cell class, as a fraction of total tissue, for each timepoint. **(D-G)** Cell type, as a fraction of each respective class, for each timepoint. Note the substantial increase in ATII-ATI transitional cells during days 3 and 7, indicating increased epithelial flux. **(H)** Fraction of each class associated with each timepoint. **(I-L)** Fraction of each cell type associated with each timepoint. Note the strong presence of cycling endothelial cells at day 7. **(M-P)** Total number of significantly perturbed ligands and receptors for each cell type at each timepoint. Note the global upregulation of cellular connectivity at day 7. Note also the bi-directional perturbation to both ligand and receptor expression in the mesothelial population at both Day 3 and Day 14 which suggests significant and complex change in mesothelial signaling behavior.

We observed an increase in immune fraction over time and a decrease in epithelial fraction (Fig 6C). We did not observe a significant change in global endothelial or mesenchymal tissue fraction during tissue expansion. Within the epithelium, ATII-ATI transitional cells notably increased at Day 3 (Fig 6D & I). The tuft and BASC fraction, conversely, remained constant, consistent with their role as stem populations supporting increased population flux. Within the mesenchyme, we observed a decrease in myofibroblasts and an increase in both mesothelial cell and pericyte fraction (Fig 6E & J). In the endothelial class, we observed a large proportion gCaps enter the cell cycle at Day 7, consistent with angiogenesis (Fig 6G & L).

These changes were associated with significant changes in the expression of cell-to-cell signaling genes (Fig 6M-P). Mesothelial cells displayed a particularly unusual pattern at day 3 (Fig 6N) and day 14 (Fig 6P), jointly elevating and reducing their expression of both ligands and receptors, meaning that they were significantly shifting their overall signaling character as both senders and receivers across multiple pathways. To probe this finding further, we performed comprehensive differential assessment of ligand-receptor connectivity over time in each sex individually, assembling a list of consensus findings showing conserved trends across datasets (see Methods). This analysis revealed 37,555 significant alterations to ligand-receptor connectivity in the tissue. Mesothelial cells were the top perturbed sending (Fig 7A) and receiving (Fig 7B) population at both Day 3 and Day 14 Feature-level investigation of signaling coming from mesothelial cells over time (Fig 7C) revealed major alterations in mesothelial expression of key morphogens during tissue regeneration. Following pneumonectomy, mesothelial cells strongly modulate their expression of *Bmp4*, a critical promoter of alveolar differentiation and homeostasis. Concurrently, they alter their expression of a wide range for stem-modulating morphogens and immuno-regulatory cytokines including, but not limited to, *Wnt4, Angptl4, Il17, Cxcl6, Il11, Fgf9, Bmp2, Wnt5a*, and *Wnt11*. These growth factors control highly disparate cellular processes in most endodermal tissues and have cell-type and context specific effects. To better understand what role mesothelial cells might be playing during tissue regeneration, we asked which cell types were most likely receiving these signals through specific receptors.

**Figure 7:**
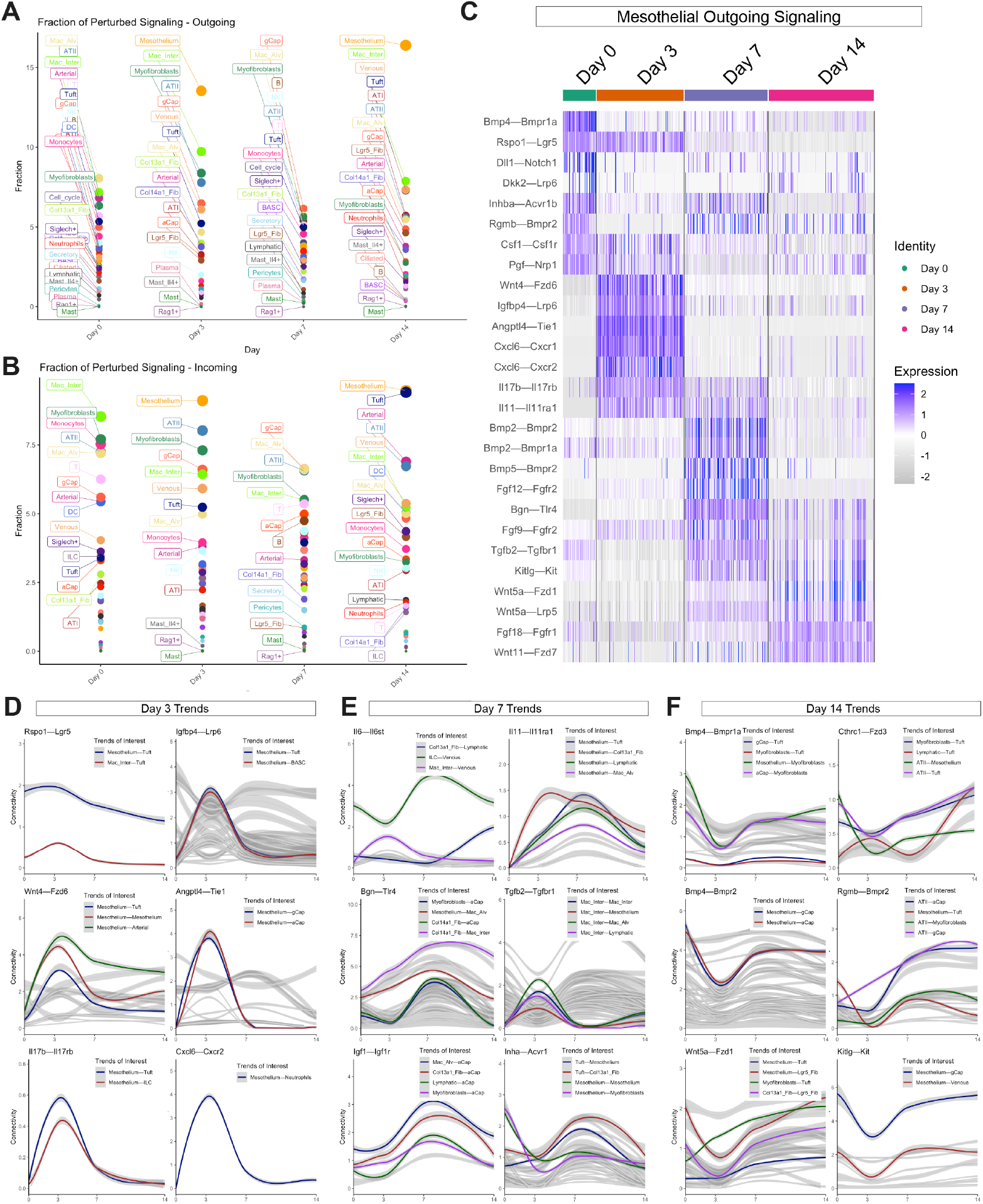
Mesothelial cells are major drivers of extracellular signaling dynamics during post-pneu-monectomy lung regeneration. **(A)** Fraction of significantly perturbed signals at each timepoint, grouped by *sending* cell type. Note that mesothelial cells are the top perturbed sending cells at both Day 3 and Day 14. **(B)** Fraction of significantly perturbed signals at each timepoint, grouped by *receiving* cell type. Mesothe-lial cells are also the top perturbed receiving cells at both Day 3 and Day 14. Note as well that tuft cells are the second most altered receivers at Day 14. **(C)** Heatmap of CellToSystem NICHES data for mesothelial cells over time. Mesothelial cells signaificantly change their signaling to other cells via WNT-, BMP-, Angpt-, Igf-, FGF-, and Cxcl-family signaling mechanisms. **(D)** Selected CellToCell trends peaking at day 3 post-pneumonectomy. At day 3, mesothelial cells drop their expression of *Rspo1*, but this change is buffered by increased tuft cell expression of *Lgr5* yielding subtly increased connectivity between these two populations. Mesothelial cells also upregulate *Wnt4*, sensed by *Fzd6* on Tuft cells and BASCs, and increase their expression of both *Igfbp4* and *Il17*, both of which activate WNT-signaling in receiving epithelial cells. We also observed factors mediating pulmonary extravasation (*Angptl4*) and immune cell chemotaxis (*Cxcl6*). **(E)** Selected signaling trends associated with day 7. Note the rise of anti-inflammatory mediators (*Il6*), matrix-mediated BMP-sensitization (*Bgn*), angiogenic cues (*Igf1*), cell proliferation factors (*Il11*), a notable reduction in *Tgfb2* signaling, and, interestingly, potential negative-feedback from Tuft cells to Mesothelial cells via *Inha*— *Acrv1*. **(F)** Selected signaling trends associated with day 14. Mesothelial cells upregulate *Bmp4* and *Wnt5a* at Day 14 which promotes epithelial differentiation. The effects of these ligands are augmented by secreted matrix proteins driving differentiation and cell polarity such as *Rgmb* and *Cthrc1*. Microvasculature-stabilizing interactions return to homeostatic levels by the end of the time-course, such as the *Kitlg*-*Kit* interaction between the Mesothelial and gCap populations. *Note that this figure represents selected mechanisms. The full 37,555 cell-to-cell signaling vectors that show meaningful differential expression during this time-course can be graphically explored via Supplements 4-6.

At day 3 (Fig 7D), mesothelial cells strongly upregulate *Wnt4*, which can be sensed by *Fzd6* on tuft cells, BASCs, and Arterial endothelium. Mesothelial cells also produce large quantities of *Igfbp4* and *Il17b*, both of which activate WNT-signaling in receiving epithelial cells (*43-45*). We found alveolar tuft cells to strongly and specifically express the receptor genes *Lrp6* and *Il17rb*, which receive each of these signals, respectively. Unexpectedly, mesothelial cells dropped their expression of the earlier-identified WNT-agonist *Rspo1* at day 3. However, the connectomic effect of this change was buffered by a concurrent increase in tuft cell expression of the cell surface receptor *Lgr5*, causing a subtle but significant increase in connectivity via this pathway stemness promoting pathway. Mesothelial cells also produced a wide variety of pro-inflammatory cues at day 3, including *Angplt4*, which targets endothelial cells and promotes extravasation (*46*), and *Cxcl2, Cxcl6, C3, Ccl7* and *Ccl20*, each of which are chemotactic for circulating immune populations (*47-51*).

Day 7 (Fig 7E) revealed an elevation of anti-inflammatory and pro-differentiation circuitry. We measured elevation of *Il10* production by interstitial macrophages and *Il6* production by innate lymphoid cells and a global drop in mesothelial production of inflammatory cytokines. We also observed notable upregulation of parenchymal, adventitial, and mesothelial secretion of matricellular factors, such as *Bgn*, which sensitize epithelial cells locally to pro-differentiation BMP-family cues (*52*). *Igf1* was elevated, which promotes microvascular sprouting angiogenesis (*53*), concurrently with *Il11*, which can be sensed by tuft cells and promotes epithelial recovery following injury (*54*). We measured both a notable reduction in *Tgfb2* signaling at day 7, and, intriguingly, a putative feedback element from Tuft cells to mesothelial cells via *Inha*—*Acrv1*, known as an extracellular modulator used by tissues to fine-tune both TGF- and BMP-family signaling gradients (*55, 56*).

By day 14 (Fig 7F), mesothelial cells upregulated *Bmp4* and *Wnt5a* back to homeostatic levels. Both of these ligands promote alveolar epithelial differentiation (*57-62*). The effects of these ligands are augmented by the earlier-produced matrix proteins driving differentiation and cell polarity, including the aforementioned *Bgn* (*52*) as well as *Rgmb (63, 64)* and *Cthrc1 (65)*. Of note, microvasculature-stabilizing interactions approach homeostatic levels by the end of the time-course, including a prominent *Kitlg*-*Kit* interaction between mesothelial and gCap populations crucial for alveolar micro-vasculature (*66*).

### Mesothelial Mechanotransduction Guides Alveolar Tuft Cell Homeostasis via WNT/BMP Balance

We next asked if mesothelial cell population displayed evidence of mechanotransduction at day 3. At day 3 in our timecourse the pleural surface would experience the greatest stress and the mesothelial matrix microenvironment would display the highest stiffness. Working from existing knowledge regarding the structure of the Hippo pathway (*67*), we created a graph-topological model of intracellular Hippo network activation state, taking into careful consideration the expected direction of transcriptomic perturbation considering mRNA flux due to pathway activation (see Methods). We then plotted each individual mesothelial cell within the resulting dimensionally-reduced space. The resulting distributions (Fig 8A) revealed a clear reduction in population-scale Hippo pathway activation in both sexes at day 3, shifting from an even On/Off spread during homeostasis (44.3% Off : 55.7% On) to a left-skewed distribution (79.9% Off : 20.1% On) at day 3. These findings directly supported mesothelial transduction of elevated tissue strain post-pneumonectomy.

**Figure 8:**
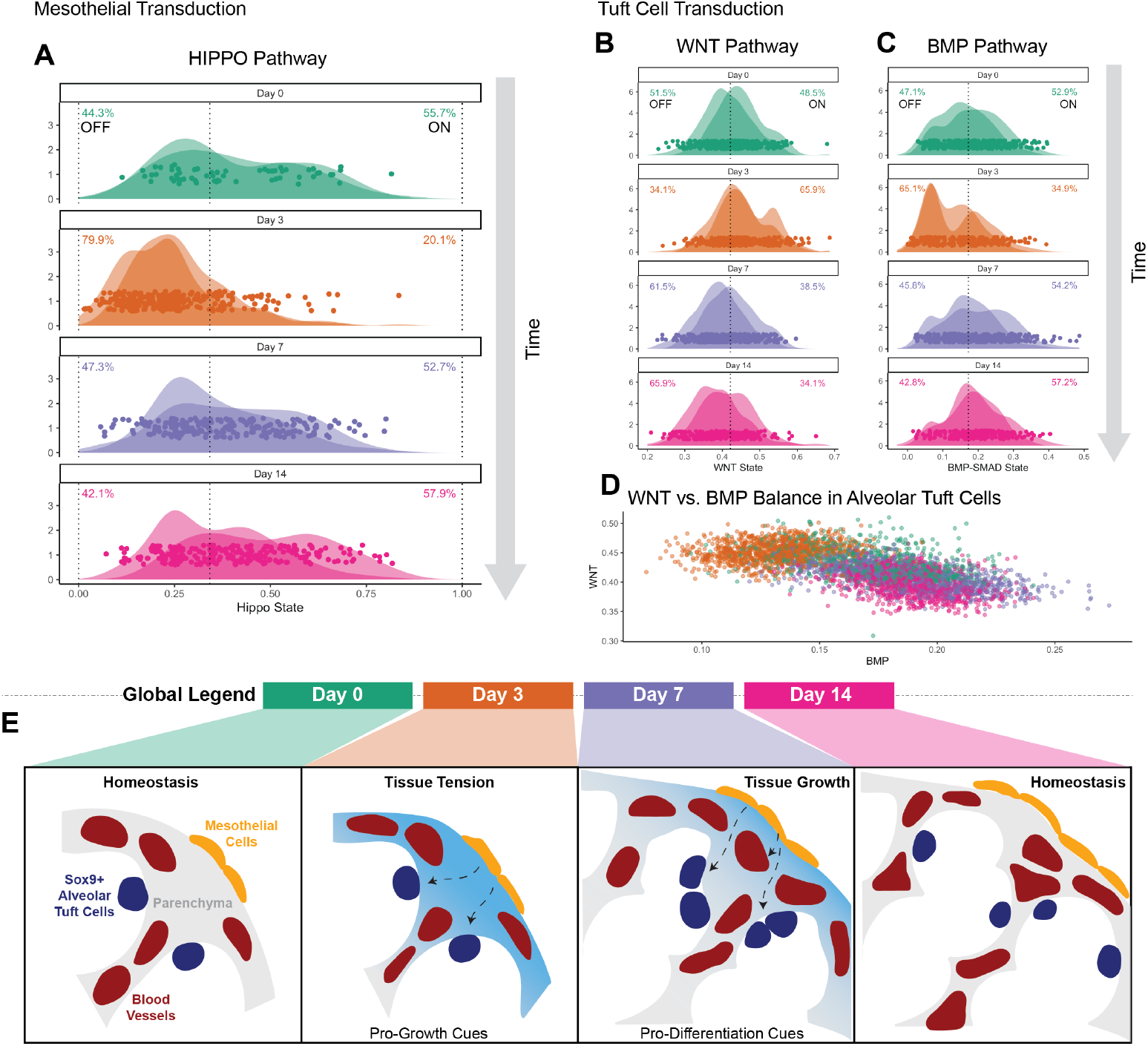
Signal transduction analysis provides evidence for sensor-effector communication between mesothelial cells and alveolar tuft cells. **(A)** Density plots of mesothelial cell HIPPO pathway state. Note the clear shift in distribution at Day 3, indicating mechanotransduction. Note that the timing of this shift correlates with the increased Mesothelial expression of WNT-family ligands and a decrease in BMP-family cues shown in Figure 7. **(B)** Density plots of tuft cell WNT-pathway state. **(C)** Density plots of tuft cell BMP-pathway state. Note the elevation of WNT-transduction and decrease of BMP-transduction at Day 3. **(D)** Scatterplot of tuft cell WNT-vs. BMP-pathway state. Note the inverse correlation. **(E)** Summary schematic. Our results support a working model in which increased pleural tension drives mesothelial cells to secrete activating growth-factors which are sensed by tissue parenchyma and alveolar-resident Sox9+/Lgr5+ Tuft cells. These cues cause the tissue to grow new parenchyma, which reduces pleural tension. As pleural tension is reduced the mesothelium produces pro-differentiation, pro-angiogenic, anti-inflammatory cues. This leads to the stabilization of a now larger organ containing new, fully vascularized, functional alveolar tissue.

We then asked if alveolar Tuft cells showed clear evidence of WNT and BMP ligand transduction. To test this, we created graph-topological models for each signal transduction network indepedently. Plotting the tuft population distribution at each time point revealed a clear elevation of tuft cells WNT transduction (Fig 8B) and decrease in tuft cell BMP transduction (Fig 8C) at day 3. Days 7 and 14, conversely, showed return of these distributions back to homeostatic patterns (Fig 8D). These findings aligned closely with our earlier evidence that tuft cell proliferation at day 7 was accompanied by high levels of mesothelial- and parenchymally-produced differentiation cues acting on receiving tuft cells. We concluded from these data that at day 3, tuft cell WNT signaling is activated and the BMP pathway is suppressed, which triggers the widespread Tuft cell proliferation that becomes clearly visibile by day 7. By day 14 these transduction trends are reversed, with WNT dormant, BMP now activated, and the Tuft population beginning to return to a quiescent, homeostatic state. These findings, in summary with the rest of our data, support a model in which alveolar tuft cells sense and respond to the mesothelial ligand secretome or to intercellular signaling cascades controlled by mesothelial mechanotransduction (Fig 8E).

## Discussion

### Summary of Findings

Our data strongly supports a regenerative function for alveolar tuft cells in the rat, identifying a powerful and underappreciated source of multipotent adult epithelial progenitors for pulmonary regenerative medicine. These cells are common in adult animals, alveolar-resident, *Sox9*- and *Lgr5*-high, WNT-responsive, expand following pneumonectomy, and are multipotent.

We also reveal Mesothelial cells as unexpected regulators of organ size and tissue growth, expressing multiple key ligands in timed coordination to regulate a controlled regenerative response. We find a sensor-effector link between mesothelium and alveolar tuft cells, as well as other alveolar parenchymal cells, based on powerful regulators including *Rspo1-Lgr5, Il17b-Il17rb, Bmp4*, and *Wnt4* vs. *Wnt5a* balance through the receptors *Fzd5* and *Fzd6*.

Mesothelium also appears to directly regulate post-pneumonectomy regenerative inflammation. After pneumonectomy, Mesothelial cells secrete large quantities of *Cxcl1, Cxcl6, C3, Ccl6, Ccl7* and *Ccl20*, all of which are chemoattractant for circulating immune cells. Mesothelium also uniquely expresses *Angptl4* post-pneumonectomy, which disrupts microvascular tight-junctions and allows immune extravasation into the alveolar space (*68*).

Return to homeostasis appears to be fine-tuned by endothelial, mesenchymal and immune effectors in combination: while Mesothelial and endothelial cells elevate their *Bmp4* expression back to normal, myofibroblast populations lay down key extracellular matrix proteins that sensitize parenchymal cells to *Bmp4* (*Bgn, (52))* and promote fibrosis-protective angiogenesis (*Nid1*, (*69, 70*)). Interstitial macrophages, meanwhile, upregulate their production of the anti-inflammatory cytokine *Il10 (71, 72)*. As pleural tension is reduced, mesothelal cells drop their production of chemoattractant cytokines and the inflammatory cascade calms down. This process is accompanied by notable elevation of mesothelial expression of *Wnt5a* (which inhibits alveolar epithelial cell proliferation), and reduces its expression of *Rspo1, Wnt4*, and *Il17b* back to homeostatic levels. The net result is new lung tissue containing functional alveolar units.

### Contextualization with Findings in Other Species

Our findings synthesize earlier work in mouse and human tissue. The alveolar tuft cells we describe here are likely functional orthologues of well-described Alveolar Epithelial Progenitors (AEPs) (*73*) in the mouse. Mouse AEPs and Rat alveolar tuft cells have similar distributions within the alveolus, show similar regenerative function, and share some gene character, with both populations displaying highly specific expression of the surface marker *Tm4sf1*.

We also identify key similarities and differences between alveolar tuft cells and bronchioalveolar stem cells (BASCs), which are known to be facultative progenitors for all pulmonary epithelial cells during homeostasis (*74-76*). BASCs are *Scgb1a1* and *Sftpc* double positive, and reside exclusively at the bronchioalveolar junction, confirmed by our data as well. Alveolar tuft cells, in the rat, reside throughout the alveoli and are negative for both *Sftpc* and *Scgb1a1*. Both BASCs and alveolar tuft cells express *Sox9, Dclk1*, and *Lgr5*, with higher levels of all three genes observed in the tuft population. Alveolar tuft cells are uniquely marked by very few genes, including *Trpm5*.

Finally, the signaling findings we have unearthed are in deep agreement with other work describing specific niche signals that regulate stem cell character in the lung. Although the observed *Rspo1*-*Lgr5 and Il17-Il17rb* circuitry from mesothelial cells to resident progenitor populations has not been previously reported, our findings do corroborate pioneering evidence from earlier research emphazing the central regulatory role of WNT-family cues within alveolar stem cell niches (*77*), the critical importance of *Bmp4* for alveolar differentiation (*57*), and the function of *Wnt5a* as a non-canonical ligand that reduces epithelial proliferation and promotes alveolar differentiation (*60, 61*).

### Contextualization with Findings in Other Organs

While BASCs have a tightly localized and well-regulated niche enviornment that reinforces multipotency during active cycling, tuft cells are able to preserve epithelial multipotency within the highly dynamic microenvironment that is the distal alveolus. We suspect this ability is deeply tied to their high *Sox9* expression, mirroring a similar pattern identifying reserve stem cells in the gut (*35*). The bronchioalveolar junction, where BASCs reside, is mechanically less variable than the cyclically stretched alveolus, and exists at a boundary between two distinct matrix and cell signaling environments. The alveolar tuft cell, in contrast, has a cell body nestled beneath ATI cells with cilia projecting into the alveolar lining layer. This means that the tuft cell is subject to both cyclic strain during breathing and a highly dynamic cell-signaling environment during normal epithelial, endothelial, and mesenchymal turnover in the alveolus. How the tuft cell maintains *Sox9* positivity in this context, in the absence of a well-defined and stable stem-cell niche, is a promising mystery. We predict that this unique ability is in part due to its unusual and highly distinct receptor profile, which both privileges the tuft cell to specific intercellular cues, in particular those produced by the mesothelium, while also isolating it from the vast majority of information flow occurring between neighboring alveolar populations.

*Sox9*, across organs, reinforces a stable, robust genetic network that sustains multipotency. Multipotency, however, is an intrinsically unstable cellular state, defined as the ability to differentially develop based on context. This means that multipotent cells have only two choices. They can either be highly susceptible to the environment and therefore only reside within a niche able to support their existence, or they can be minimally susceptible to their environment but maintain a tightly regulated internal network state that maintains their multipotent character, as if in hibernation. The *Sox9*+ BASCs and tuft cells in the rat lung display exactly this binary pattern: one (the BASC) exists in a tightly regulated environment and displays active multipotency (associated with low *Sox9*-positivity), creating progeny continually that differentiate based on their developing context. The other (the tuft cell) exists in a highly dynamic environment, where it is well positioned to sense necessary cues to expand or adapt, while maintaining a highly quiescent (*Sox9*-high), relatively inflexible cell state that maintains and protects multipotency until it is directly told to activate by tissue-specific sensor cells.

### The Lung is an Ecosystem Regulated by Sensor-Effector Communication

In order to activate latent multipotency, proliferate, and differentiate, cells must respond to their microenvironments. To do so, they leverage threshold-like genetic switch mechanisms responsive to transduction pathways driven by physiologic variables. This general biological pattern synthesizes the above findings and helps to explain the evidence of focused communication between mesothelial cells and alveolar tuft cells. The data presented here is consistent with alveolar tuft cells playing the role of sentinel-like reserve stem cells within the tissue: they are primed to receive highly specific morphogenic cues from sensor cells within the lung. Their high Sox9 level simultaneously makes them multipotent (flexible in response direction) as well as robust to perturbation (difficult to influence without the right cues, quiescent). These properties allow them to lie in waiting until they receive the appropriate signals to activate.

We note from our experiments that elevation of WNT-signaling from mesothelial cells may induce tuft cells to exit their quiescent state and to enter an actively proliferating stem-like state. Mesothelial cells, which reside only on the surface of the lung, are in a uniquely priviledged to sense tissue strain. In combination with their unique ligand profile, they are a prime candidate for further investigations on the basic mechanisms of lung regeneration and pathology in a variety of contexts. The data we have presented here requires a great deal more exploration, as well as mechanistic knock-down studies to more deeply investigate the role of specific factors mediating mesothelial-parenchymal communication and control. Genetic tools to perform such experiments, unfortunately, are not well developed in the rat, suggesting that investment may be required in this basic technology if we are to better understand this phenomenon.

We conclude by noting that BASCs and tuft cells together represent a powerful cellular population for pulmonary bioengineering. Adult stem cells are crucial tools for regenerative medicine, because they provide a way for us to create new tissues directly from patient samples without having to induce pluripotency and then reverse-engineer cellular reprogramming. The discovery of a robust, reliable adult progenitor population that can be manipulated *in vitro* and is clearly dynamic *in vivo* provides a much needed scientific foothold. We hope that the reported findings might be leveraged to engineer advanced therapeutics and to build a foundation for a unified consensus vision of how tissue homeostasis is regulated across multiple organs.

## Supporting information

Supplement 1 - Key Resources Table

## Acknowledgements

We would like to thank Dr. Ruslan Medzhitov, Dr. Themis Kyriakides, Dr. Yuval Kluger, and Dr. Aaron Osgood-Zimmerman for their support and counsel during this work and the preparation of the manuscript.

We would like to thank Dr. Lisa Leffert, Dr. Helene Benveniste, and Dr. Robert Schonberger as well for their continual mentoring and encouragement during the startup of this laboratory. We would also like to thank Dr. Carlos Fernandez-Hernando and the Vascular Biology and Therapeutics Program at Yale for their continuing and essential support.

We are grateful to Dr. Guilin Wang and Dr. Christopher Castaldi in the Yale Center for Genome Analysis for their helpful support and patience throughout the many years of this project.

We would like to thank Dr. Scott Magness for generously discussing his work with us and providing much-needed encouragement that we were studying something valuable and not an artifact. We would also like to thank him for sending Sox9-GFP mouse lung tissue for histologic analysis. The work on this study was supported in part by grant T32GM086287 from the National Institute of General Medical Sciences (NIGMS). The opinions expressed are those of the authors and do not necessarily represent the thoughts or opinions of NIGMS, NIH, or the United States government. This work was supported by laboratory startup funding from the Department of Anesthesiology within the Yale School of Medicine.

Additional funding was provided by Else Kröner-Fresenius Foundation (EKFS) grants EKFS 2021_EKEA.16 and 2020_EKSP.78 (J.C.S.); CORE100Pilot (Advanced) Clinician Scientist Program of Hannover Medical School funded by EKFS and the Niedersächsisches Ministerium für Wissenschaft und Kultur, and the German Research Foundation (SCHU 3147/4-1) (J.C.S.)

This paper was typeset with the bioRxiv word template by @Chrelli: www.github.com/chrelli/bioRxiv-word-template

## Author Contributions

Conceptualization, TO, MSBR, LEN, AL, NK;

Methodology, TO, SM, TA, JY, KL, AMG;

Software, JY, JCS, MSBR;

Validation, RR;

Formal Analysis, TO, TA, JY, HK, MSBR;

Investigation, TO, SM, SE, RR, HK, TA, JCS, AMG, YY, MSBR;

Resources, NW, KL;

Data Curation, TO, SM, NW, HK, AG, SE, MSBR;

Writing – Original Draft, TO & MSBR;

Writing – Review & Editing, AG, TT, TO, SM, MSBR, AL, LEN, NK;

Visualization, TO, JY, TA, AMG, AL, MSBR;

Supervision MSBR, DS, AL, LEN, NK;

Funding Acquisition TT, DS, JCS, MSBR, LEN, NK.

## Data Availability

Raw FASTQ files, extracted digital gene expression matrices, and processed R objects containing all relevant metadata are available at Gene Expression Omnibus GEO ******.

Scripts used for this project are publicly available on GitHub at https://github.com/RaredonLab/Obata2023.

Supplements are available on FigShare at https://figshare.com/projects/Obata2023/191505.

## Materials and Methods

### In vivo

#### Animal husbandry

All rats were wild-type Sprague Dawley rats (Rattus norvegicus) obtained from Charles River (8-10 weeks old, 300-350 g) and bred and maintained at the animal center of Yale University. All experiments were performed under the NIH Guidelines for the Care and Use of Laboratory Animals and approved by the Yale University Committee on Animal care and use.

#### Rat lung dissociation

Rats were selected between 7-9 weeks old, or from selected experimental conditions described below, and were anesthetized with ketamine 75 mg/kg and xylazine 5 mg/kg and injected IP with 400 U/kg sodium heparin. The abdomen was opened in sterile fashion and the xiphoid process lifted to reveal the underside of the diaphragm. The diaphragm was punctured and retracted to reveal the chest cavity. The rib cage was lifted and an incision was made axially to reveal the trachea. The trachea was separated from the esophagus, opened using fine scissors, cannulated with a barbed fitting, and secured with 4-0 silk. The right ventricle was opened with scissors, and the pulmonary artery (PA) cannulated with an 18-gauge feeding needle and secured with 4-0 monofilament polypropylene. An incision was made in the left ventricle and the lungs were perfused via the PA with 100 mL of PBS containing 100 U/mL sodium heparin and 0.01 mg/mL sodium nitroprusside with concurrent airway ventilation. The heart-lung block was then removed from the chest to be perfused with enzyme.

DMEM containing 1mg/mL Collagenase/Dispase (Roche), 3 U/mL Elastase (Worthington), and 20 U/mL DNAse (Worthington) pre-heated to 37C was used to dissociate all tissues with the exception of a single sample, rat811, which was dissociated with Liberase TM (Sigma). Enzyme was perfused through the vasculature and then instilled into the trachea 5-10x. Care was taken to fully recruit and inflate the alveolar regions of the lung and to have the tissue fully saturated with enzyme. Following inflation and/or perfusion with enzyme, non-lung tissue was removed, including the heart and extralobular airways. The remaining tissue and effluent were placed on a rocker at 37°C and 60rpm for 20 minutes. At this point the lung tissue was pushed through a fine wire sieve using a weighing spatula until only dense connective tissue remained. The strainer was then rinsed with 20 mL ice-cold DMEM containing 10% FBS, 1% penicillin/streptomycin, 1% amphotericin, and 0.1% gentamicin. The resulting tissue solution was spun down for 5min at 300g, the supernatant discarded, and the pellet resuspended in red cell lysis buffer at a 1:1 ratio with pellet volume. After 120 seconds at room temperature, the buffer was diluted with 0.01% BSA in PBS (0.1mg/mL) and spun down a second time. The pellet was resuspended in 0.01% BSA in PBS and filtered through a 70μm filter. The cells were then spun for 3 min at 300g, resuspended in 0.01% BSA in PBS, and passed twice through a 40μm filter. The resulting cell suspension was counted, assessed for viability, and serially diluted to the desired concentration for single cell library preparation or Dclk1+ cell isolation.

#### Dclk1+ cell isolation

DCLK-1-positive cells were isolated from 220-250 g Sprague-Dawley rats (approximately 6–8-week-old) using magnetic beads selection. Briefly, rats were anesthetized, the thorax was exposed, and the lungs were cannulated. All blood was flushed and extracted *en bloc*, as described for rat lung dissociation. The lungs were perfused through the pulmonary artery and inflated via the airway with DMEM containing 1 mg/ml Collagenase/Dispase (Roche), 3 U/ml Elastase (Worthington), and DNAse I (Worthington). Each lobe to be used was then carefully dissected and placed in a conical tube with dissociation buffer, followed by incubation for 20 min in a shaking water bath at 37°C. During incubation, the lung tissue becomes soft and friable, allowing it to be directly pushed through a sterile stainless-steel sieve with a weighing spatula which acts to mechanically dissociate the tissue in a low-shear setting. Ice-cold DMEM containing 10% FBS was used to rinse the sieve and quench the enzymes. Larger collagenous structures not naturally passing through the sieve were discarded. The below cell-matrix suspension was collected in a conical tube and centrifuged at 300 g for 5 min at 4°C. The supernatant, containing suspended matrix, was discarded. The cell pellet was suspended in red cell lysis buffer (Gibco) and incubated for 2 min at room temperature, before being diluted and spun down again. The supernatant was discarded and the cell pellet resuspended in 0.4 mg/ml BSA. This suspension was sequentially passed through 70- and 40-μm cell strainers and incubated with anti-DCAMKL-1 antibody (Abcam) conjugated magnetic beads with polyclonal sheep anti-rabbit IgG (Invitrogen) for 30 min at 4°C. After repeated rinses, the cells were sorted using the Dynamag (Invitrogen) protocol, and freshly isolated DCLK-1 cells were used for further culture or analysis.

#### Pneumonectomy

Male Sprague–Dawley rats aged 8–12 weeks were selected for pneumonectomy. Rats were anesthetized with ketamine (75 mg/kg) and xylazine (5 mg/kg), orally intubated with a 16G catheter, and maintained using isoflurane with an animal ventilator during the procedure. All procedures were performed under aseptic conditions. The left chest was shaved, disinfected with ethanol and iodine, and immobilized in the right lateral position on the operating table. Thoracotomy was performed using a left intercostal incision just caudal to the left scapula (at 4-5 intercostal space). All membranous structures attached to the left lobe were dissected, leaving only the pulmonary hilum attached, and the pulmonary artery/vein and the left main bronchus were then collectively ligated using 7-0 polypropylene thread. After detaching the hilum, the entire left lobe was removed from the thoracic cavity and discarded. The wound was sutured closed, and the rat was withdrawn from anesthesia and awakened. Post-surgical pain control was provided per Yale IACUC guidelines.

#### 5-ethynyl-2’-deoxyuridine (EdU) Pulse-Chase Experiments

EdU was injected intraperitoneally into the rats at 50 mg/kg 24 hours before planned sacrifice. For sacrifice, the rats were anesthetized and cannulation was performed as described above for dissociation. Heparin saline was used to flush blood from the lungs until the tissues were completely white. The lungs were then inflated, via the trachea, with optimum cutting temperature (OCT) compound until full recruitment was obtained. The bronchus was then ligated and the tissue removed and frozen in a disposable mold containing OCT. 6 μm lung sections were cut using a cryotome, air dried at -20C, and stored until further use. Upon removal of the frozen sections from the freezer, the tissue sections were immediately fixed using 50 μl of 4% formaldehyde in PBS (pH7.4) for 15 min at room temperature. The lung sections were then covered with PBS containing 3% BSA for 5 min, and then rinsed three times with PBS. Cells were permeableized by covering the sections with 0.5% Triton X-100 in PBS for 20 min before being rinsed 3 times in PBS for 5 min each. The sections were then incubated in blocking buffer containing 5% serum from the secondary antibody’s host in PBS, pH 7.5, for 30 min to block unspecific binding of the antibodies. Alternative blocking buffers are 1% BSA or 1% FBS in PBS. EdU was labeled using the in vivo imaging kit (BCK-EdU594, base click, Saint-Aubin, France) according to the manufacturer’s instructions. Double immunostaining was then performed against target cell markers.

### In vitro

#### Organoid culture

8–10-week-old rats were used to generate lung organoids. Briefly, freshly sorted lineage-labeled Dclk1+ cells were resuspended in LPM-3D medium (LPM-3) and mixed with growth factor-reduced Matrigel (BD Biosciences) in a 1:1 ratio; 90 μL of the mixture was placed in the 0.4-μm pore Transwell inserts (Corning) placed in 24-well plates (Corning) using pre-chilled pipette tips. Organoid mixtures were incubated at 37°C in 7% CO2/air for 30 minutes to allow mixture to solidify. 500 μL of LPM-3D medium was then placed below the Transwell insert. In some experiments, sorted fibroblasts were seeded into 24-well plates (Corning), below the Transwell insert, and expanded for 7-14 days for further organoid co-culture assays with Dclk1+ cells. For monocultures, 1x10^5^ Dclk1+ cells were seeded in each insert. For co-cultures, approximately 0.5-1 × 10^4^ Dclk1+ cells and 0.5-1 × 10^5^ fibroblasts were seeded in each insert. 500 μL of LPM-3D medium was placed in the lower chamber. All components of LPM-3D medium can be referenced in supplementary table (S_).

#### Organoid dissociation

Organoids were dissociated for single-cell analysis and cryopreservation using an adapted protocol from Ali et al., 2020 (*78*). LPM-3D culture medium was aspirated and 1 mL of ice-cold DPBS was added to each insert. The organoid mixture was then gently pipetted up and down with large-bore tips to dissociate the matrigel. Samples of the same condition were pooled into a single conical and centrifuged at 300g for 5 minutes at 4°C. 0.5 mL of TrypLE Express (Gibco, 12605010) was added to the suspension and incubated at 37°C for 10-15 minutes. Following incubation, the suspension was gently pipetted up and down ∼10 times to obtain a single cell suspension. The suspension was then centrifuged for 10 minutes at 4°C and 300 g to form a pellet. The supernatant was removed and 10 mL of Advanced DMEM/F12 medium was added to stop TrypLE Express activity. Suspension was again centrifuged at 300 g for 5 minutes at 4°C to obtain a pellet. Cells for cryopreservation were resuspended in freezing medium containing 90% FBS and 10% DMSO (w/ 1% PenStrep) and snap frozen. Cells for single-cell library preparation were resuspended in a 0.4 mg/ml BSA solution at 1x10^*6*^ cells/ml and sent for library preparation.

#### Single-cell library preparation and sequencing

Single cell RNA sequencing libraries were generated with the Chromium Single Cell 3’ v2 and v3 assays (10X Genomics) as well as an in-house constructed DropSeq platform (*79*). Samples were diluted for an expected cell recovery population of 10,000 cells per lane per sample. Libraries were sequenced on the Hiseq4000 platform (Illumina) to a target depth of 50,000 reads per cell.

#### Rat lung fixation

Rat lungs were surgically isolated as above, cleared of blood, inflated with approximately 5mls of air (half inflation) using a 10ml syringe, and then perfused intravascularly via the PA with either room-temperature 10% neutral-buffered formalin or 4% paraformaldehyde warmed to 37C. Perfusion of fixative was done either via gravity at 30 cmH_*2*_O with a re-circulation pump (if a partial lung block) or via direct perfusion at 50ml/min (for a full set of lungs) with a pulse-dampener between the pump and the tissue. Perfusion of fixative was continued for a minimum of 30 minutes. The tissue was then transferred into a beaker of fixative, incubated for 4 hours at room temperature, and then transitioned to 70% ethanol for storage and down-stream processing.

#### Immunohistochemistry on paraffin sections

Samples of native rat lung and organoids were fixed in formalin for 3-4 hours, then paraffin-embedded and sectioned at Yale Pathology Tissue Services (YPTS). Hematoxylin and eosin (H&E) staining of sections was carried out by YPTS or in-house. To localize protein expression in paraffin-embedded samples, sections were de-waxed in xylene and ethanol, followed by soaking in citric acid buffer (0.01M citric acid, 0.05% Tween 20, distilled water, pH 6.0) at 75°C for 20 min. After cooling at room temperature, all sections were rinsed in PBS and permeabilized for 15 min in permeabilization solution (0.2% Triton X-100 in PBS)for 15min. Sections were blocked (5% BSA and 0.75% glycine in PBS), and incubate for overnight at 4°C with primary antibodies (see list of antibodies used in Supplement 1: Key Resources Table). Antibodies were diluted in blocking buffer to the appropriate concentration. Primary antibodies were removed and slides were rinsed in PBS. Following rinses, sections were incubated in Alexa Fluor conjugated secondary antibodies at 1:500 dilution for 1 hour at room temperature. Sections were stained with DAPI for 1min and mounted with polyvinyl alcohol with DABCO (PVA-DABCO).

#### Immunofluorescence on frozen sections

Tissue designated for frozen sections was fixed with formalin and embedded in optimal cutting temperature (OCT) gel. The embedded tissue was frozen over dry ice for ∼25 min., then transferred to the -80C freezer until sectioned. To stain sections for, sections were thawed to room temperature and rehydrated in PBS for 10 min. Antigen retrieval was performed by treating the sections with 6M urea in glycine-HCl for 10 min at room temperature (RT), per methods described by Hinenoya et al. (*80*).

Sections were blocked with 10% normal goat serum for 15 min at RT and primary antibodies to collagen IV alpha 2 and alpha 5 chains were applied in PBS w/ 1 mg/mL bovine serum albumin (BSA) for 60 min RT at a 1:50 dilution. Sections were washed with PBS and incubated with secondaries in blocking buffer for 30 min at RT at 1:100 dilution. Tissue was then washed a final time with PBS, incubated with DAPI and mounted with PVA-DABCO.

#### Stitched-field fluorescence microscopy

Whole-lung stitched fluorescence images were obtained using an EVOS FL Auto Imaging System (ThermoFisher Scientific). In brief, multi-stained fluorescence slides were placed on the microscope, and the light intensity and exposure were adjusted appropriately for each channel. The entire tissue was then imaged in the DAPI channel, and the imaging range was selected. Channel settings were optimized for this selected imaging range and saved. The entire imaging range was then automatically scanned, with autofocusing, according to the saved channel settings. The obtained tiled images were reconstructed as one large, stitched image, which was used for further analysis and image quantification.

### In silico

#### Single-cell data extraction and filtration

Data was aligned and extracted using STARsolo with the default parameters for each chemistry (*81*). The top 30,000 cell barcodes per sample, ranked by nUMI, were then imported into R for further processing. Ensembl Rnor6.0 release 92 was used to align the data.

#### Single-cell data cleaning, clustering, and annotation

The data for each sample was first cleaned (conservatively) individually, following the excellent guidelines presented by (*82*). Cleaned samples were then grouped together and cleaned further, if necessary, through iterative clustering and removal of multiplet and low-information clusters. One a fully cleaned dataset was obtained, cells were ‘classed’ into either epithelium, endothelium, immune, or mesenchymal categories. Each category was then subclustered and annotated independently.

#### Statistical analysis of single-cell data

We were interested in focusing our analysis on genes that were specifically and strongly up- or down-regulated across the post-pneumonectomy time course in *both* the male and female data. We therefore constructed our significance testing strategy to reflect this scientific goal. First, we split the dataset into two categories, male and female. Then, we split each of those datasets by cell type annotation, yielding a total of 34x2=68 distinct R objects. Differential expression, by timepoint, was then run for each of these objects using the FindAllMarkers function in Seurat. Minimal thresholds were used for this first, global test: we did not limit output in any way except to require a minimum of 3 cells per feature and minimum of 3 cells per group (test.use = ‘wilcox’, min.cells.feature = 3, min.cells.group=3, min.pct = 0, logfc.threshold = 0, return.thresh=1, only.pos = FALSE). The output from this approach yielded 68 output lists describing the significance of differential expression for nearly all captured genes, in all cell types, at each point in the time course.

We then assembled a ‘consensus’ data frame combining the results from each of these sex-specific studies. First, we assembled two large data frames, each capturing significance testing results from each sex. We annotated each data frame with additional metadata describing unique row identifiers (gene-CellType-Timepoint combinations) and added columns designating if a given row was found to be up or down regulated in a given sex. We then combined information from the two sex-specific data frames into a single, consensus data frame, preserving all information captured from both sexes. We computed mean log fold changes and combined p-values using Fisher’s Method. Reported significant findings had to display directional consensus between the male and female datasets and had to have a combined adjusted p-value of less than 0.05.

#### Connectomic analysis

Analysis of ligand-receptor signaling was performed using NICHES v1.0.0 (*37*). Data was imputed with ALRA (*83*) prior to running NICHES. Imputation was performed for the entire dataset at once and was limited to genes expressed in greater than 50 cells in each object.

Our significance testing strategy for these connectomic data mirrored that described above. First, we split the dataset into two categories, male and female. Then, we split each of those datasets by VectorType annotation (CellType-CellType crosses), yielding a total of 34x34x2=2312 distinct R objects. Differential expression, by timepoint, was then run for each of these objects using the FindAllMarkers function in Seurat. Minimal thresholds were used for this global connectomic test as well: we did not limit output except to require a minimum of 3 cells per feature and minimum of 3 cells per group (test.use = ‘wilcox’, min.cells.feature = 3, min.cells.group=3, min.pct = 0, logfc.threshold = 0, return.thresh=1, only.pos = FALSE). The output from this approach yielded 2312 output lists describing the significance of differential connectivity for nearly all queried ligand-receptor mechanisms, in all cell type crosses, at each point in the time course. Reported significant connectomic findings had to display directional consensus between the male and female datasets, had to have a combined adjusted p-value of less than 0.05, and had to display trend-agreement with the raw data on either the sending or the receiving side of the connectomic interaction.

Seurat v4.3.0 (*84*), ggplot2, and ComplexHeatmap v2.14.0 (*85*) were used to plot selected significant findings, trends, and patterns of interest.

#### Intracellular pathway analysis

Pathway-specific gene modules were defined based on literature review. Every gene was then assigned either a 1 or a -1, based on the expected direction of change in the context of elevated pathway transduction. This assignment had to be done carefully, based on the known role of each protein within a signaling network, and whether we would therefore expect increased or decreased *transcription* of the corresponding gene, which is what our data represents. These assignments allowed us to compute a comparable activation score, for every queried cell, of the defined pathway. We have chosen to visualize these data using 1-dimensional density plots. The values on the x-axis have been unity normalized so that a value of zero represents the maximum ‘off-ness’ of a pathway and a value of 1 represents the maximum ‘on-ness’ of that same pathway, given our measured data range.

#### Quantitative image analysis in R

Images were processed using EBImage (*86*). Briefly, the data for each channel was loaded, normalized, and processed using adaptive threholding. Data tables were then created quantifying pixel-positivity across each channel. The resulting data were then plotted using ggplot2.

**Figure S1:**
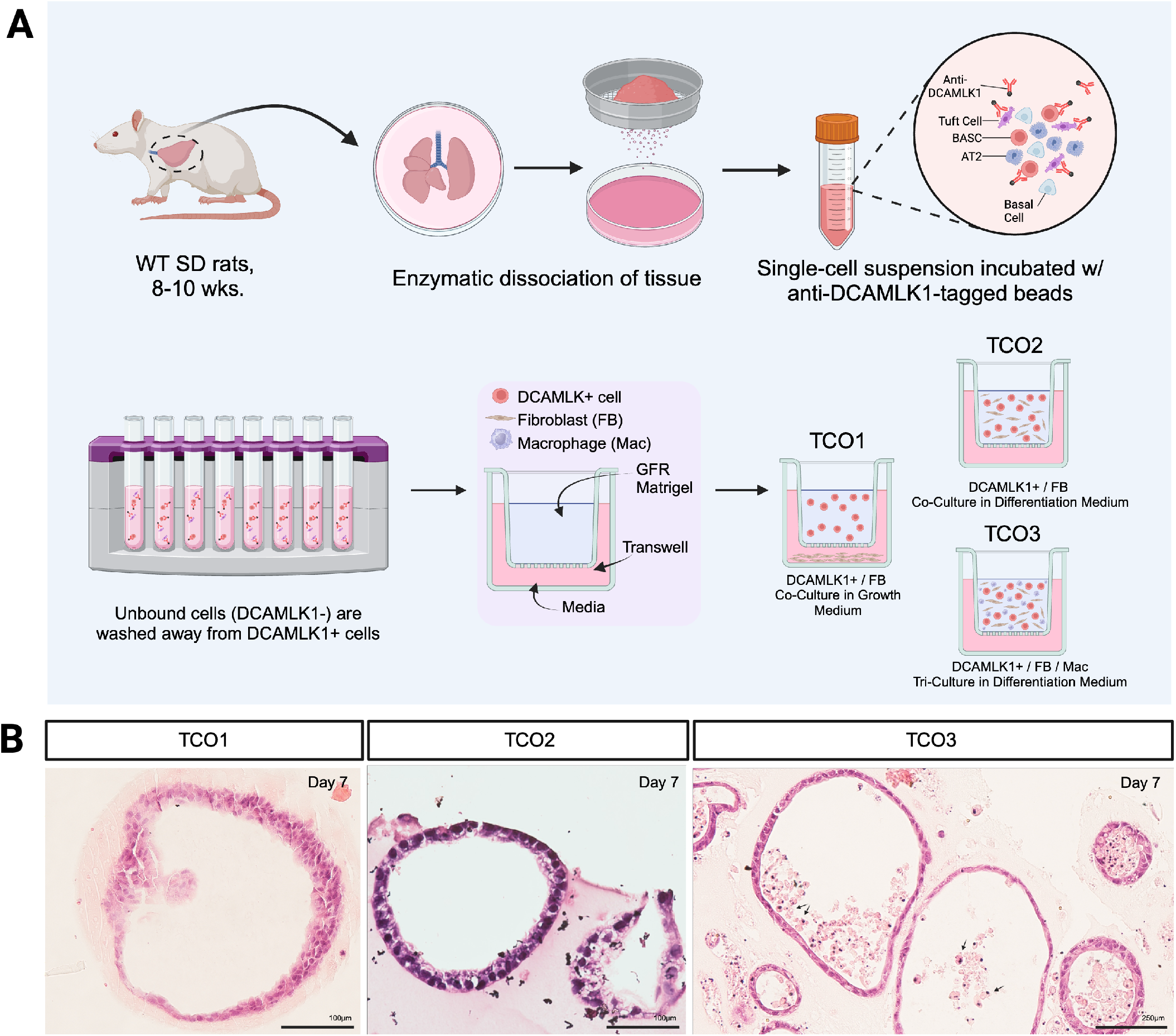
Organoid Experimental Design

## List of Supplements

Supplement 1: Key Resources Table - FigShare

Supplement 2: Adult Rat Atlas – GEO ******

Supplement 3: Post-Pneumonectomy Time Course Atlas – GEO ******

Supplement 4: Niche-wise Significant Findings - FigShare

Supplement 5: Influence-wise Significant Findings - FigShare

Supplement 6: Mechanism-wise Significant Findings - FigShare

Scripts used for all data processing in this project have been compiled on GitHub at https://github.com/Ra-redonLab/Obata2023

